# MHA, an interactive website for scRNA-seq data of male genitourinary development and disease

**DOI:** 10.1101/2022.07.29.501961

**Authors:** LiangYu Zhao, YiFan Zhao, ChenCheng Yao, Zheng Li, YingBo Dai, YuXin Tang

**Affiliations:** Department of Urology, Department of Interventional Medicine, Guangdong Provincial Key Laboratory of Biomedical Imaging, The Fifth Affiliated Hospital, Sun Yat-sen University, Zhuhai, Guangdong Province 519000, China; Department of Andrology, the Center for Men’s Health, Urologic Medical Center, Shanghai Key Laboratory of Reproductive Medicine, Shanghai General Hospital, Shanghai Jiao Tong University School of Medicine, Shanghai 200080, China; The Sinotech Genomics CO. LTD. Shanghai 200080, China

## Abstract

The development of single-cell sequencing technology has expanded the understanding of cell heterogeneity and disease progression in the male genitourinary system. However, complex processing processes and unprofessional analytical annotations limit the daily use and wildly sharing of published datasets. Here, we describe the Male Health Atlas (MHA), an easy-to-operate online visual website focused on the male genitourinary system, allowing researchers to quickly search for expression patterns of one or multiple genes, and dynamic changes of gene expression during tissue development or under pathological states. At present, the MHA includes nine objects, two species, five types of male genitourinary tissues/organs, and 258,777 single-cell profiles. Compared with other single-cell databases, the MHA has the following advantages: (1) It is easy to operate, and users can complete retrieval and output of results within 5 min without any bioinformatics skills; (2) The selection and analysis of data are under professional supervision of researchers in the male genitourinary field; (3) It contains most of the main male-specific organs and will continue to be updated; (4) MHA provides multi-dimensional output patterns, including tSNE/UMAP, violin, and bubble plot output for displaying single or multiple genes. MHA is the first single-cell database website in the field of andrology and male reproduction, providing researchers with scRNA-seq resources and an accessible tool. MHA is freely available at http://malehealthatlas.cn/.

## Introduction

The development of single-cell RNA (scRNA) sequencing technology makes it possible to study the heterogeneity of tissue and organ microenvironments at molecular and single-cell levels [1]. At present, scRNA data for almost all tissues and organs of human and animal models (mostly mice) have been published [2–4]. However, in-depth mining of these data often requires bioinformatics skills, and the tedious operation limits the ability of researchers to quickly retrieve the required information. From the perspective of daily use, what researchers typically need is the expression characteristics of a specific target gene during a physiological process or disease progression, retrieved within a few minutes through simple tab clicking and input.

Some comprehensive scRNA databases have not focused on the male genitourinary system and are prone to problems, such as irregular disease classification and cell clustering [5]. For example, spermatogenesis is a complex and continuous cellular developmental process, which comprises mitotic spermatogonia, meiotic spermatocytes, and post-meiotic spermiogenesis from round spermatids to spermatozoa, and spans dozens of cell subtypes [6]. The early stages of meiosis alone can be divided into at least four stages, however, comprehensive databases often simply define meiosis as one to three germ cell clusters [5, 7]. Researchers in reproductive medicine often cannot gain useful information from such rudimentary analysis. Another example, in human corpus cavernosum tissue, endothelial and smooth muscle cells show functional and transcriptional differences according to whether they belong to cavernosal trabecular or nearby vessels. Such important information is often overlooked, except by andrologists or related researchers. To solve the problems mentioned above, we present the Male Health Atlas (MHA), an easy-to-operate online visual website with curated scRNA datasets of male genitourinary systems to better assist researchers in this field.

## DATA COLLECTION AND CONTENT

### Data collection

The first version of MHA contains nine objects, two species (Homo sapiens and Mus musculus), five organs/tissues (testis, epididymis, vas deferens, corpus cavernosum, and prostate), eight cell types, and 258,777 unique scRNA data profiles. These data were derived from published studies or our unpublished scRNA data.

- **Human_Testis_Development_Atlas:** ten human testicular samples were from 2- (n=1), 5- (n=1), 8- (n=1), 11- (n=1), and 17-years-old (n=1), and adult (n=5) subjects, to show the normal developmental process of human testes. This dataset comes from our previous published study [8].
- **Mouse_Testis_Development_Atlas:** nine testicular samples were from 3- (n=1), 6- (n=1), 8- (n=1), 11- (n=1), 14- (n=1), 17- (n=1), 21- (n=1), 25- (n=1), and 35-day-old (n=1) C57-mice, to show the normal developmental process of mouse testes. This dataset comes from our unpublished scRNA data.
- **Human_Germ_Cell_Lineage_Atlas:** this dataset is the germ cell line subset of the “Human_Testis_Development_Atlas”, from spermatogonial stem cells to spermatids, in which we provide simple and detailed clustering classifications.
- **Mouse_Germ_Cell_Lineage_Atlas:** this dataset is the germ cell line subset of the “Mouse_Testis_Development_Atlas”, from spermatogonial stem cells to spermatids.
- **Human_Testis_Non_Obstructive_Azoospermia_Atlas:** this dataset included five adult (23–31 years-old) testicular samples with normal spermatogenesis and seven testicular samples from non-obstructive azoospermia (NOA) patients (26–32 years-old). NOA can be further classified as idiopathic nonobstructive azoospermia (n=3), Klinefelter’s syndrome (n=3), and YqAZFa microdeletion (n=1) according to etiology. This dataset comes from our previous published study [8].
- **Human_Epididymis_Atlas:** this dataset contains three caput epididymis samples from normal adults aged 31, 57, and 32 years [9].
- **Mouse_Epididymis_Vas_deferens_Atlas:** this dataset contains samples of caput, cauda, corpus, and vas deferens from mice. Each population in this dataset represented a pool of eight tissue samples obtained from four male mice sacrificed at 10–12 weeks of age [10].
- **Human_Corpus_Cavernosum_Atlas:** eight samples of corpus cavernosum from normal men (n=3) and patients with erectile dysfunction (ED, n=5). Patients with ED can be further classified as non-diabetic ED (non-DM, n=3) and diabetic ED (DMED, n=2). We provide simple and detailed clustering classifications. This dataset comes from our previous published study[11].
- **Human_Prostate_Cancer_Atlas:** this dataset contains 16 human prostate samples from three young male organ donors aged 18–31 years [12] and 13 prostate cancer samples from patients aged 61–81 years (12 primary and 1 lymph node metastasis) [13].

### Processing and annotation

The raw input files for each dataset have been processed (cell-gene matrix) with CellRanger. Because the cell type and origin of different tissues vary greatly, different quality control criteria were used. For example, we set “nFeature_RNA < 9000 & percent.mt < 40 & nCount_RNA <80000” for testicular cells and “nFeature_RNA < 6000 & percent.mt < 15 & nCount_RNA <10000” for prostate cells. The Seurat R package was used for cell clustering and dimensionality reduction processing. Each dataset was annotated according to the age, disease, and tissue origin of the samples.

Because researchers often have different requirements (more or less detailed) for cell annotation in different situations, we provide at least two different levels of cell classification methods for users to choose (named as “cell_type” and “cell_type2”). For example, in the annotation of “cell_type” of the “Human_Germ_Cell_Lineage_Atlas” dataset, the entire germ cell lineage was divided into 14 stages, and spermatocytes were divided into five stages. However, sometimes users do not need such detailed classification criteria. Therefore, for the annotation of “cell_type2”, we combined some stages and left only four stages of spermatogenesis. This annotation convention applies to most datasets in the MHA.

### Website construction

To better embed related functions from the Seurat R package for single-cell data visualization, such as feature, violin, and dot plots, the whole MHA website has been constructed using R shiny package. Based on the current architecture of MHA, it usually takes half a minute to load a dataset. To ensure fast access, the website is deployed on an Alibaba Cloud cluster, with 4 cores and 64 GB of RAM.

## DATA ACCESS

The MHA website is available online at http://malehealthatlas.cn/ and requires no registration. At the output end, we provide UMAP and TSNE, violin, frequency bar, and bubble plots to meet various data display requirements of users. The MHA has been tested using Microsoft Edge, Google Chrome, and Apple Safari browsers. All image results are free to download by clicking the “Download” button (higher resolution), and can also be saved by “right-click” and “save as” (lower resolution). The website also provides the differentially expressed genes (DEGs) list between groups or cell types in each dataset.

## RESULTS

### MHA statistics

Nine datasets of the MHA cover two species, five organs/tissues, 57 samples, eight cell types, and 258,777 unique scRNA datasets (Table 1 and Figure 1A-I). Among them, 71% and 29% of cells were obtained from human and mouse samples, respectively (Figure 1J), and 68% of cells were obtained from healthy tissues and 32% of cells belonged to tissues under pathological states (Figure 1K). Except for seminal vesicles and ejaculatory ducts, most of the major male reproductive organs have been included, such as testis (63%), epididymis (4.1%), vas deferens (1.1%), corpus cavernosum (25%), and prostate (6.5%) (Figure 1L). Eight major cell types include the epithelial cells (Epithelial, 8.05%), endothelial cells (EC, 10.94%), Schwann cells (SWC, 0.05%), Sertoli cells (SC, 9.94%), muscle cells (Muscle, 13.18%), Leydig cells (LC, 16.68%), immune cells (Immune, 4.6%), germ cells (Germ, 24.72%), and fibroblasts (FB, 11.83%) (Figure 1M).

**Table 1.** Basic information of the nine datasets.

| Projects | Organism | Tissue/Organ | State | No. of samples | No. of cells |
| --- | --- | --- | --- | --- | --- |
| Human_Testis_Development_Atlas | Homo sapiens | Testis | Normal; Growth and development | 10 | 61605 |
| Mouse_Testis_Development_Atlas | Mus musculus | Testis | Normal; Growth and development | 9 | 66442 |
| Human_Germ_Cell_Lineage_Atlas | Homo sapiens | Testicular germ cells | Normal; Growth and development | 10 | 22777 |
| Mouse_Germ_Cell_Lineage_Atlas | Mus musculus | Testicular germ cells | Normal; Growth and development | 9 | 25802 |
| Human_Testis_Non_Obstructive_Azoospermia_Atlas | Homo sapiens | Testis | Normal and Disease | 12 | 62515 |
| Human_Epididymis_Atlas | Homo sapiens | Epididymis | Normal | 3 | 5299 |
| Mouse_Epididymis_Vas_deferens_Atlas | Mus musculus | Epididymis and Vas deferens | Normal | 4 | 8880 |
| Human_Corpus_Cavernosum_Atlas | Homo sapiens | Corpus cavernosum | Normal and Disease | 8 | 64740 |
| Human_Prostate_Cancer_Atlas | Homo sapiens | Prostate | Normal and Disease | 16 | 16893 |
| Total (Duplication between projects were eliminated ) |  |  |  | 81 (57) | 334953 (258777) |

**Figure 1.**
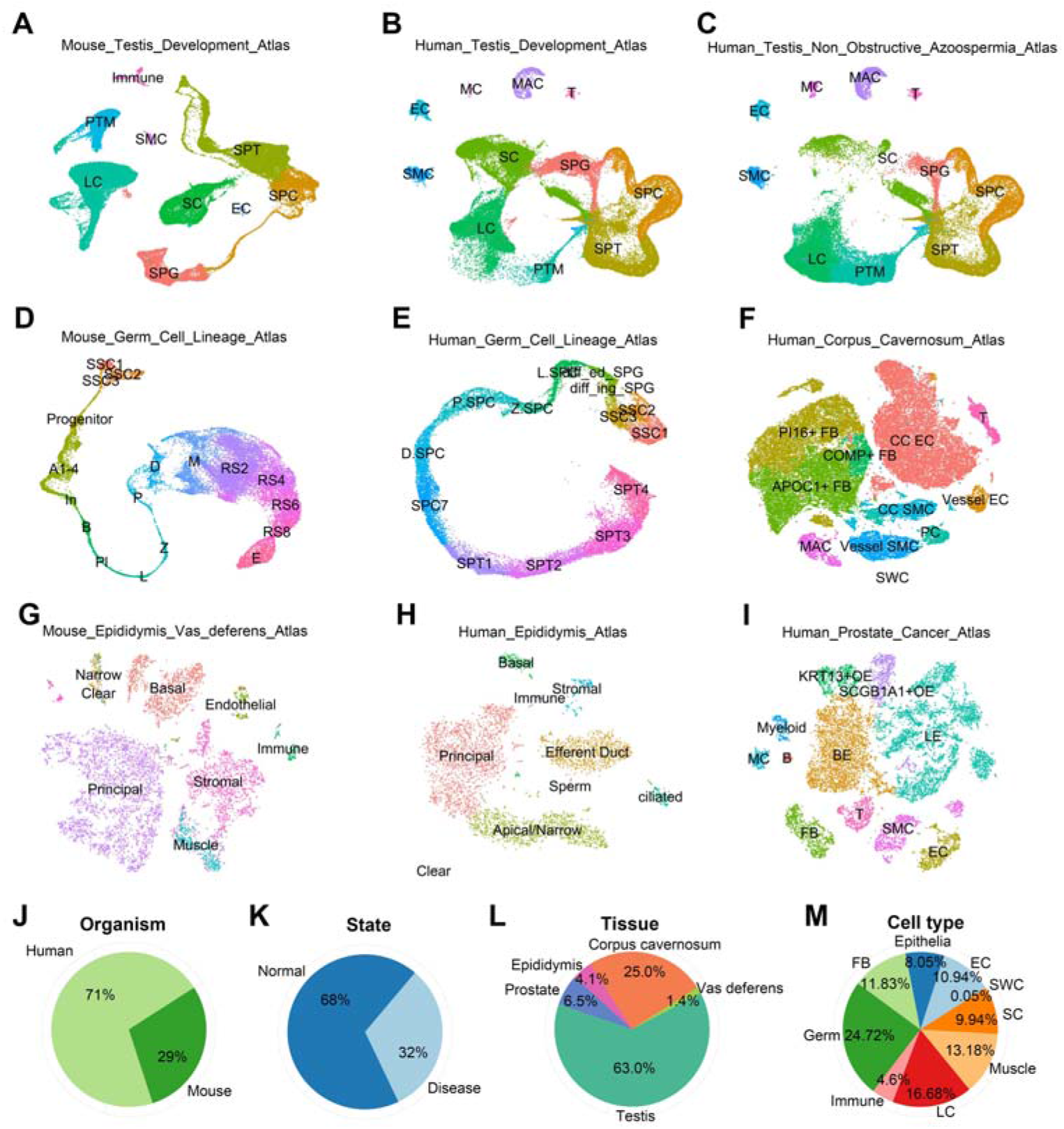
Male Health Atlas (MHA) statistics information. (A–I) The t-distributed stochastic neighbor embedding (tSNE) or uniform manifold approximation and projection (UMAP) plots show the cell composition of each dataset. Cells are colored according to their “cell_type” annotation. (J–M) The statistics information of all nine datasets of MHA, including the ratio of cells belonging to human or mouse samples (J), cells under normal or disease state (K), tissue sources of cells (L), and cell types (M). EC, endothelial cells; FB, fibroblasts; LC, Leydig cells; SC, Sertoli cells; SWC, Schwann cells.

### Web interface

The MHA provides a public user interface to easily and quickly collect characteristics of gene expression. On the “HOME” page, users can browse the introduction and dataset information to quickly understand the goals and functions of MHA. This page will also list news about the latest research progress of our team and the citation format for MHA. The “DATASETS” page is the functional page of the MHA, where users can select data sets to be displayed visually. In addition, users can find the instructions of MHA in the “ABOUT” page and contact information in the “AUTHORS” page (Figure 2A).

**Figure 2.**
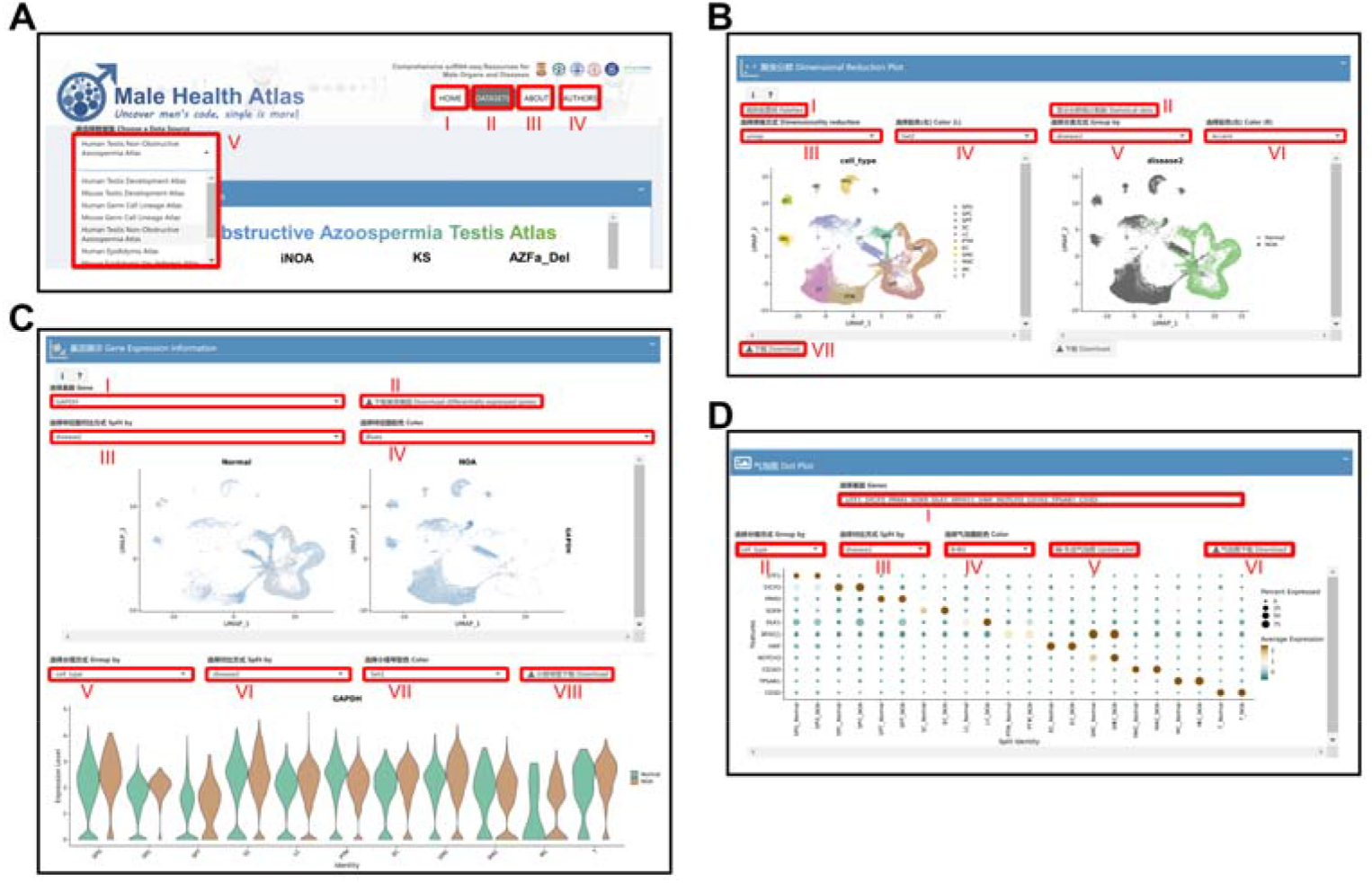
The main interface page of Male Health Atlas (MHA). (A) The main pages and dataset of MHA. (I) The HOME page introduces the MHA database, updates, and references; (II) the “DATASETS” page shows output results; (III) the “ABOUT” page shows help and manual instructions for using MHA; (IV) the “AUTHORS” page shows information on the authors of MHA. (B) The dimensional reduction plot of the chosen dataset. (I) Shows all color schemes that could be used; (II) shows the ratio of cell types; (III) selects the dimensional reduction option; (IV) selects the color scheme of the left panel; (V) selects the annotation option of the right panel; (VI) selects the color scheme of the right panel. (C) Single gene expression display of the chosen dataset. (I) Allows input or selection of a specific gene; (II) downloads the DEGs list of this dataset; (III) selects a split method for dimension reduction plot; (IV) selects the color scheme to show different expression levels of the gene; (V) selects a grouping method for the violin plot (factors of X-axis); (VI) selects a split method within one group; (VII) selects the color scheme of the violin plot; (VIII) downloads the violin plot results. (D) Display of the expression of multiple genes of the chosen dataset. (I) Allows input or selection of one or more genes; (II) selects a grouping method for the dot plot; (III) selects a split method within one group; (IV) selects the color scheme of the dot plot to show different expression levels of genes; (V) creates or refreshes the dot plot; (VI) downloads the dot plot results.

After entering the “DATASETS” page, users can click “Palettes” to view all potential color schemes of the output results. The ratio of each cell cluster or group can be viewed by clicking “Statistical data”. In addition, umap and tSNE plots will be provided to show the cell annotation of each dataset. Users can easily change the cell annotation and color schemes (Figure 2B).

In the “Gene Expression Information” section, umap or tSNE plots, and violin plots will be provided for users to obtain information of single gene expression. Users can easily change the cell annotation and color schemes according to their requirements (Figure 2C).

A “Dot plot” section is provided for information about the expression levels of multiple genes. Users can input multiple genes and change the cell annotation and color schemes according to their requirements. The size of dots represents the ratio of positive to negative cells and the color scale bar represents the expression level (Figure 2D).

## DISCUSSION

### Advantages of MHA

The physiological processes and diseases of the male genitourinary system represent an active research area. As of July 7, 2022, there were 453,813 results obtained by searching the keyword “male genitourinary system” in PubMed. In recent years, increasing numbers of studies have used scRNA-seq technology in this field to provide new insight in male-specific tissues at the molecular and single cell levels. However, the depth of data mining and convenience of operation for this technology are often contradictory for subsequent single cell data analysis. In most cases, researchers only want to know the patterns or levels of gene expression for selective genes, as well as their dynamic changes during development and disease. In short, researchers need a fast, convenient, and professional interactive platform to display processed scRNA-seq data.

Although some analysis platforms for scRNA-seq can provide functions such as data expression, pseudo-time analysis, and cell interaction, they are often complicated to operate [14]. In addition, many scRNA databases contain millions of cells and all major tissues and organs, but their cell annotation methods are not focused on the male genitourinary system and are often not monitored by trained investigators in andrology or urology [2, 4, 7]. To our knowledge, there is no convenient tool for the targeted analysis of scRNA-seq data for the male genitourinary system. Therefore, we established the MHA to enable researchers in our field to quickly access gene expression information of major published scRNA datasets, as well as for crossdisciplinary researchers to accurately obtain relevant annotation results.

### Future perspectives

The MHA covers the physiological development and common diseases of the male genitourinary system based on nine scRNA-seq datasets. In the future, the MHA will be improved in three directions. First, a separate page will be developed for animal models, including gene knockout, mutant alleles, and other artificial disease models. For example, as far as we know, there have been at least a dozen scRNA-seq results reported for the testes of knockout mouse models. For the testis, we will combine single-cell data of gene knockout or mutant mouse models with the data of normal mice to form a new dataset, which will help researchers to quickly understand the effect of a specific gene on the testicular microenvironment and spermatogenesis. Secondly, the timeline dataset will be expanded (especially for the testis), covering from embryo development to aging, to form a single-cell roadmap of the whole lifecycle of the male genitourinary system. Third, the MHA will contain information from more species. Some studies, such as erectile dysfunction models, typically use rats, so scRNA-seq data from the cavernosa of the rat penis will be essential for such studies. In addition, we will continue to update the database and user experience once a year around December.

## ACKNOWLEDGMENTS

This work was supported by the China Postdoctoral Science Foundation (2021M703747, L.Z.), GuangDong Basic and Applied Basic Research Foundation (2021A1515111109, L.Z.), National Natural Science Foundation of China (82171597, Z.L.; 81871215, Z.L.; and 82071636, Y.T.), and open funds of Guangdong Provincial Key Lab of Biomedical Imaging (GPKLBI202104, L.Z.). We thank Charles Allan, PhD, from Liwen Bianji (Edanz) (www.liwenbianji.cn/) for editing the English text of a draft of this manuscript.

## REFERENCES

1. Wen L, Tang F. Recent advances in single-cell sequencing technologies. Precision clinical medicine. 2022;5(1):pbac002. Epub 2022/07/14. doi: 10.1093/pcmedi/pbac002. PubMed PMID: 35821681; PubMed Central PMCID: PMCPMC9267251.

2. Jones RC, Karkanias J, Krasnow MA, Pisco AO, Quake SR, Salzman J, et al. The Tabula Sapiens: A multiple-organ, single-cell transcriptomic atlas of humans. Science (New York, NY). 2022;376(6594):eabl4896. Epub 2022/05/14. doi: 10.1126/science.abl4896. PubMed PMID: 35549404.

3. Tabula Muris C, Overall c, Logistical c, Organ c, processing, Library p, et al. Single-cell transcriptomics of 20 mouse organs creates a Tabula Muris. Nature. 2018;562(7727):367–72. Epub 2018/10/05. doi: 10.1038/s41586-018-0590-4. PubMed PMID: 30283141; PubMed Central PMCID: PMCPMC6642641.

4. Han X, Wang R, Zhou Y, Fei L, Sun H, Lai S, et al. Mapping the Mouse Cell Atlas by Microwell-Seq. Cell. 2018;172(5):1091–107.e17. Epub 2018/02/24. doi: 10.1016/j.cell.2018.02.001. PubMed PMID: 29474909.

5. Qu J, Yang F, Zhu T, Wang Y, Fang W, Ding Y, et al. A reference single-cell regulomic and transcriptomic map of cynomolgus monkeys. Nature communications. 2022;13(1):4069. Epub 2022/07/14. doi: 10.1038/s41467-022-31770-x. PubMed PMID: 35831300; PubMed Central PMCID: PMCPMC9279386.

6. Zhao L, Zhu Z, Yao C, Huang Y, Zhi E, Chen H, et al. VEGFC/VEGFR3 Signaling Regulates Mouse Spermatogonial Cell Proliferation via the Activation of AKT/MAPK and Cyclin D1 Pathway and Mediates the Apoptosis by affecting Caspase 3/9 and Bcl-2. Cell Cycle. 2018;17(2):225–39. Epub 2017/11/25. doi: 10.1080/15384101.2017.1407891. PubMed PMID: 29169284; PubMed Central PMCID: PMCPMC5884123.

7. Han L, Wei X, Liu C, Volpe G, Zhuang Z, Zou X, et al. Cell transcriptomic atlas of the non-human primate Macaca fascicularis. Nature. 2022;604(7907):723–31. Epub 2022/04/15. doi: 10.1038/s41586-022-04587-3. PubMed PMID: 35418686.

8. Zhao L, Yao C, Xing X, Jing T, Li P, Zhu Z, et al. Single-cell analysis of developing and azoospermia human testicles reveals central role of Sertoli cells. Nature communications. 2020;11(1). doi: 10.1038/s41467-020-19414-4.

9. Leir SH, Yin S, Kerschner JL, Cosme W, Harris A. An atlas of human proximal epididymis reveals cell-specific functions and distinct roles for CFTR. Life science alliance. 2020;3(11). Epub 2020/08/29. doi: 10.26508/lsa.202000744. PubMed PMID: 32855272; PubMed Central PMCID: PMCPMC7471510.

10. Rinaldi VD, Donnard E, Gellatly K, Rasmussen M, Kucukural A, Yukselen O, et al. An atlas of cell types in the mouse epididymis and vas deferens. Elife. 2020;9. Epub 2020/07/31. doi: 10.7554/eLife.55474. PubMed PMID: 32729827; PubMed Central PMCID: PMCPMC7426093.

11. Zhao L, Han S, Su H, Li J, Zhi E, Li P, et al. Single-cell transcriptome atlas of the human corpus cavernosum. Nature communications. 2022;13(1):4302. Epub 2022/07/26. doi: 10.1038/s41467-022-31950-9. PubMed PMID: 35879305.

12. Henry GH, Malewska A, Joseph DB, Malladi VS, Lee J, Torrealba J, et al. A Cellular Anatomy of the Normal Adult Human Prostate and Prostatic Urethra. Cell Rep. 2018;25(12):3530–42.e5. Epub 2018/12/20. doi: 10.1016/j.celrep.2018.11.086. PubMed PMID: 30566875; PubMed Central PMCID: PMCPMC6411034.

13. Chen S, Zhu G, Yang Y, Wang F, Xiao YT, Zhang N, et al. Single-cell analysis reveals transcriptomic remodellings in distinct cell types that contribute to human prostate cancer progression. Nature cell biology. 2021;23(1):87–98. Epub 2021/01/10. doi: 10.1038/s41556-020-00613-6. PubMed PMID: 33420488.

14. Jiang A, Lehnert K, You L, Snell RG. ICARUS, an interactive web server for single cell RNA-seq analysis. Nucleic acids research. 2022. Epub 2022/05/11. doi: 10.1093/nar/gkac322. PubMed PMID: 35536286; PubMed Central PMCID: PMCPMC9252722.

